# Bayesian rhythmic model for jointly detecting circadian biomarkers and predicting molecular circadian time in human post-mortem brain transcriptome

**DOI:** 10.64898/2026.01.21.700905

**Authors:** Xiangning Xue, Hao Wang, Zihan Zhu, Guan Yu, Marianne L. Seney, Liangliang Zhang, George C. Tseng

## Abstract

Transcriptomic circadian analysis of human post-mortem brain provides a unique opportunity to characterize in vivo molecular circadian rhythms across brain regions implicated in aging and psychiatric disorders. A primary goal in such analyses is the detection of circadian biomarkers. However, this task is complicated by the frequent mismatch between a subject’s recorded circadian clock time and their true molecular circadian time — arising from observational or recording errors, as well as intrinsic biological variability. Existing methods typically address either biomarker detection or circadian time prediction in isolation. Because errors in one task can degrade performance in the other, the lack of a unified approach remains a key limitation. We propose BayCT — a Bayesian model for simultaneous circadian marker detection and molecular circadian time estimation. The model extends naturally to repeated measurements from multiple brain regions or organs. For circular data, we employ a von Mises prior distribution, with slice sampling and reversible-jump Markov chain Monte Carlo (MCMC) for Bayesian inference. Through extensive simulations and applications to transcriptomic data from three human brain regions and from 12 mouse organs, BayCT demonstrates superior performance in both biomarker detection and circadian time estimation. Furthermore, we highlight the advantages of integrating data across brain regions, achieving substantial improvements in both tasks.

## Introduction

Circadian rhythms are endogenous ∼ 24-hour oscillations that synchronize physiological activities to the daily light–dark cycle. This intrinsic timekeeping system optimizes the timing of diverse biological processes, thereby enhancing organismal survival. At the molecular level, circadian rhythms are driven by a transcriptional–translational feedback loop that regulates thousands of circadian-controlled genes in a coordinated manner (Pittendrigh, 1960). In mammals, the suprachiasmatic nucleus (SCN) serves as the master clock, maintaining self-sustained oscillations independent of environmental cues (Aschoff, 1960). Although the free-running period is close to — but not exactly — 24 hours, it varies with age, sex, and ancestry (Crowley and Eastman, 2018; Eastman et al., 2015), necessitating entrainment to environmental signals such as light to maintain precise phase alignment. However, modern lifestyle factors — including artificial lighting, altered work schedules, and late-night blue light exposure — challenge this entrainment process (Finger and Kramer, 2021; Tähkämö, Partonen and Pesonen, 2019). Chronic circadian misalignment, such as that induced by rotating night shifts, has been linked to elevated risks for cancer and other chronic diseases (Schernhammer et al., 2001; Wegrzyn et al., 2017).

Detection of circadian biomarkers from high-throughput omics data is fundamental for studying rhythmic regulation in health and disease. A variety of computational methods have been proposed, including parametric cosinor regression (Cornelissen, 2014), non-parametric approaches such as JTK_CYCLE (Hughes, Hogenesch and Kornacker, 2010), spectral methods such as the Lomb–Scargle periodogram (Glynn, Chen and Mushegian, 2006), and deep learning methods such as BIO_CYCLE (Agostinelli et al., 2016). Flexible models like JTK_CYCLE and BIO_CYCLE can capture diverse waveform shapes beyond the sinusoidal assumption of cosinor regression. However, they may suffer from inflated or deflated type I error rates and present challenges in accurately estimating rhythmic parameters such as phase and amplitude (Ding et al., 2021).

Our motivating example is from a human post-mortem brain transcriptomic study for circadian research, in which bulk RNA-seq data are profiled in three brain regions - nucleus accumbens (NAc), caudate and putamen - of the same *N* = 59 subjects with time of death (TOD) information for each subject (Section 4.2). In post-mortem human studies, a subject’s recorded time of death (TOD) often deviates from their true molecular circadian time (MCT), due to observational or recording errors and intrinsic inter-individual variation in circadian phase. Such variability arises from environmental influences (e.g., light exposure, temperature), behavioral patterns (e.g., sleep–wake schedules, meal timing), genetic differences (Jones et al., 2019), and lifestyle-induced chronodisruption. Accurate estimation of MCT has substantial implications for chronotherapy, which aims to align treatment timing with an individual’s circadian phase to maximize efficacy and minimize toxicity (Dallmann, Okyar and Lévi, 2016; Cederroth et al., 2019). Meta-analyses and clinical trials have shown that dosing time can significantly influence drug toxicity and therapeutic response (Printezi et al., 2022; Ruben et al., 2019).

Several computational approaches have been developed to predict MCT from transcriptomic data. Early work such as the Molecular Timetable (Ueda et al., 2004) correlated samples to reference gene-phase profiles, while more recent methods employ machine learning (e.g., ZeitZeiger (Butte and Hastie, 2016), TimeSignature (Braun et al., 2018)) or Bayesian inference (e.g., Tempo (Auerbach et al., 2022) and CHIRAL (Talamanca, Gobet and Naef, 2023)). Deep learning models such as CYCLOPS (Anafi et al., 2017) and Cyclum (Liang et al., 2020) leverage nonlinear representation learning to improve prediction accuracy. However, these methods address MCT prediction and circadian biomarker detection as separate tasks, despite the fact that errors in one can propagate and degrade performance in the other. Figure 1A visually demonstrates the need for joint modeling of both tasks by scatter plots for time of death (TOD; x-axis) versus gene expression level (y-axis) in four selected core clock genes (CIART, CRY1, PER2 and PER3) in 59 human caudate samples. Subject 103 shows clear deviance between recorded TOD (solid circle) and the predicted MCT (empty circle). Correcting recorded TOD=-5.61 to predicted MCT=0.43 yields improved sinusoidal fits for circadian biomarker detection, and these gains in biomarker detection subsequently enhance the accuracy of MCT prediction.

**Fig 1.**
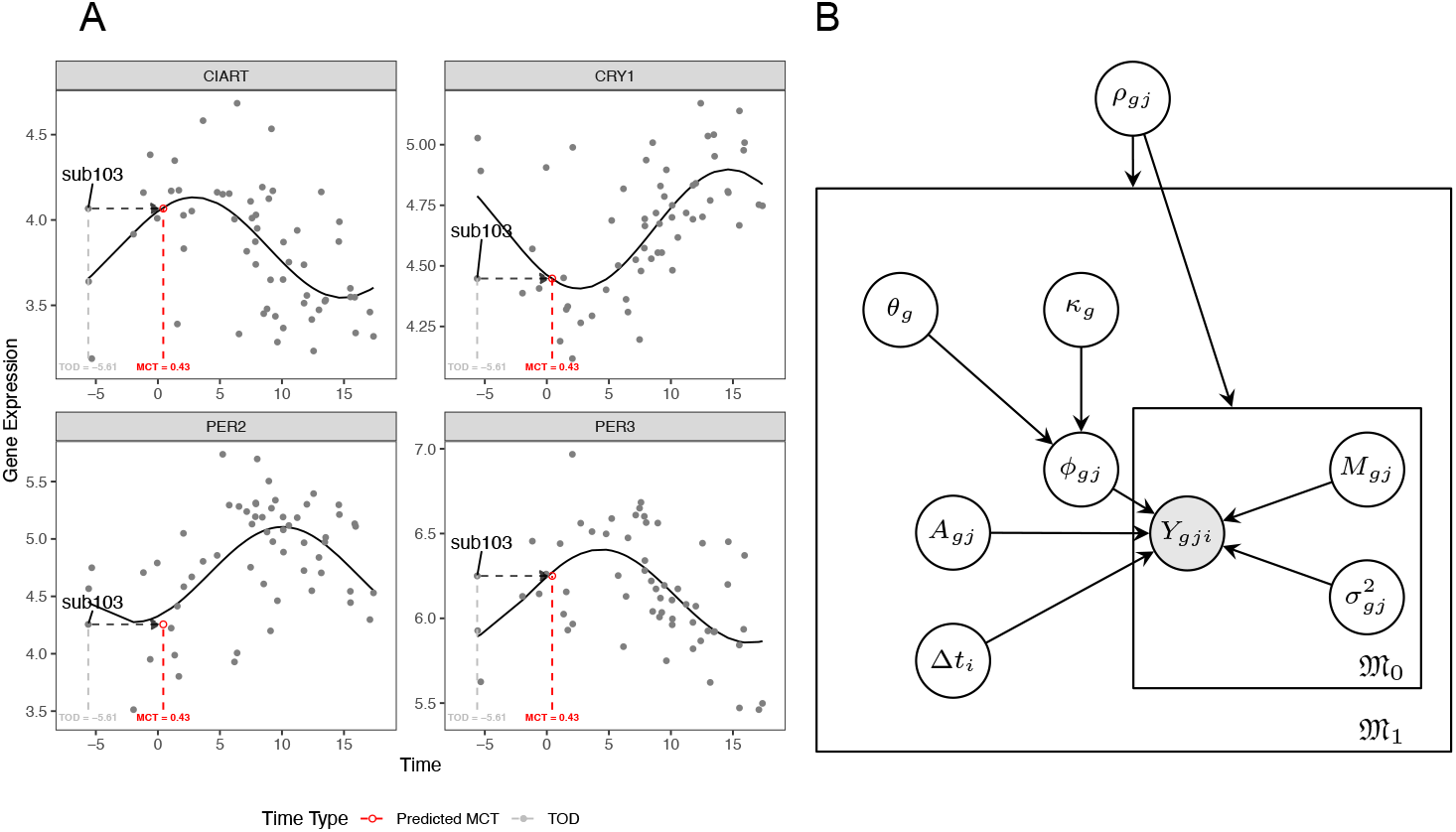
A Circadian gene expression patterns of four selected core clock genes in the human caudate. x axis is time on a ZT scale (-6 to 18 hour) and y axis is gene expression level. The black line is the fitted sinusoidal curve using TOD. Correcting TOD=-5.61 to predicted MCT=0.43 yields improved sinusoidal fits for circadian biomarker detection, and the improvement in biomarker detection will subsequently enhance MCT prediction accuracy. B. Graphical representation of BayCT-M. The graphical representation of BayCT is a simplified version without subscript for brain region j and parameters θ_g_ and κ_g_.

To address this gap, we propose BayCT — a unified Bayesian model for simultaneous Circadian biomarker detection and molecular circadian Time estimation in Section 2.1. BayCT employs a von Mises prior to accommodate the circular nature of time and uses slice sampling and reversible-jump Markov chain Monte Carlo (MCMC) for efficient Bayesian inference (Section 2.2). In Section 2.3, we further extend the model to BayCT-M to integrate data from multiple brain regions or organs. In Section 3 and Section 4, we evaluate BayCT and BayCT-M through extensive simulations and real-data applications involving transcriptomic measurements from 12 mouse organs and three human brain regions. Our results demonstrate substantial improvements in both biomarker detection and MCT estimation, and highlight the benefits of integrating data across multiple brain regions or organs.

## 2. Methods: BayCT and BayCT-M

In this section, we present the BayCT and BayCT-M model. In Section 2.1, we begin with introducing a basic Bayesian circadian model (BayC) without aligning molecular circadian time, followed by a full BayCT model including MCT estimation. The BayC model will also be used as a baseline method in numerical studies when MCT correction is ignored in biomarker detection. Section 2.2 shows the corresponding Monte-Carlo Markov Chain (MCMC) sampling procedures for Bayesian inference. Finally, we extend the BayCT model to BayCT-M (Section 2.3) allowing multi-brain-region integrative analysis.

### 2.1. BayCT model

Consider *Y*_*ig*_ the expression values of *G* genes (1 ≤*g* ≤*G*) and *N* subjects (1 ≤*i* ≤*N*) with the time of death (TOD) of subject *i* standardized to zeitgeber time (ZT) as *t*_*i*_. Here, zeitgeber time is adjusted by factors such as time zone, daylight savings time, and sunrise time (calculated by date of death and longitude and latitude of the place of death) and and ZT=0 is standardized to 6:00AM as previously described (Chen et al., 2016). For simplicity, we begin with a basic Bayesian model for circadian marker detection without TOD correction by MCT, which is denoted as BayC model. The cosinor model of BayC for a given gene *g* is:

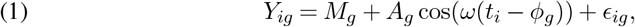

where *A*_*g*_ is the oscillation amplitude, *ϕ*_*g*_ is the peak time of gene expression, *M*_*g*_ represents the MESOR (midline-estimating statistic of rhythm; mean expression level), *ϵ*_*ig*_ is Gaussian noise with 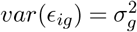, and 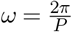 is the oscillation frequency (*P* = 24 hours is pre-defined to reflect circadian rhythm). The range of the amplitude and the peak time are constrained with *ϕ* ∈ [0, 24] and *A* ∈ (0, +∞).

The hyperparameters and priors are set as:

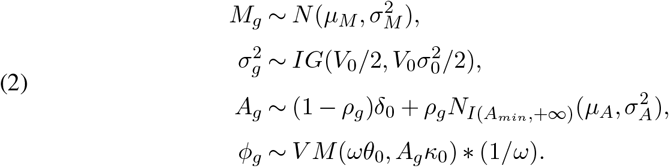

Here, *N* is normal distribution, *IG* is inverse gamma, *A*_*g*_ follows a spike-and-slab prior with spike at 0 and slab as truncated normal distribution *N*_*I*(*a,b*)_ constrained within (a,b), *V M* is von Mises distribution for circular data. The indicator variable *ρ*_*g*_ = 1 indicates gene *g* is rhythmic and *ρ*_*g*_ = 0 arryhythmic. We set 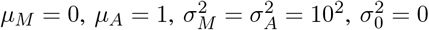 and *V*_0_ = 2 to give minimally-informative prior for *M* and *σ*. For prior of *A, A*_*min*_ can be user-defined, representing the smallest biologically meaningful amplitude but we use *A*_*min*_ = 0 throughout the paper for simplicity. The von Mises prior for *ϕ* is used for its periodic nature. We set *θ*_0_ = 0 and *κ*_0_ = 0 to give non-informative prior to *ϕ*. In fact, *V M* (*θ*_0_, 0) is equivalent to Uniform(0, *P*).

We note that the value of the circadian indicator *ρ* generates two possible model states for the gene expression below with different parameter space: Θ_0_ *=*{*M* ^′ 2′^} *ρ* = 0for and Θ_1_ = *M, A, ϕ, σ*^2^ for *ρ* = 1, which calls for reversible jump sampling to be introduced in Section 2.2.

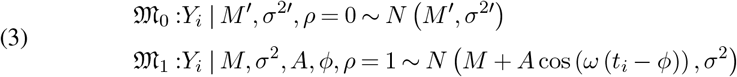

To achieve joint biomarker detection and time correction in BayCT model, we introduce 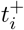 to denote the underlying true molecular circadian time (MCT). The circadian time deviation between MCT and TOD is denoted as Δ*t*_*i*_ such that 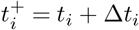. The cosinor model in BayCT simply replaces *t*_*i*_ in the BayC model (Equation 1) with 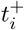:

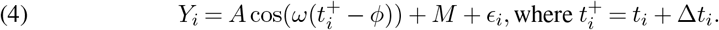

Since Δ*t*_*i*_ is also periodic, we give a reasonably informative prior

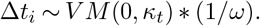

The mean of the prior is set to 0, and *κ*_*t*_ is set to 6, which is equivalent to a standard deviation of about 1.5 hours. Figure 1B shows the graphical representation of BayCT under 𝔐_0_ and 𝔐_1_.

### 2.2. Sampling procedure

The sampling procedure for the BayCT model consists of two alternating stages: updating within models and transitioning between the two models in Equation 3 using reversible jump MCMC. The complete procedure is divided into four steps. Step I occurs only when updating within model 𝔐_0_, while Steps I, II, and IV are applied to model 𝔐_1_. Step III is performed after each update within either model.

Step I updates the two parameters *M* and *σ*^2^ shared by 𝔐_0_ and 𝔐_1_. In Step II, the application of slice sampling for jointly updating *A* and *ϕ* in 𝔐_1_ significantly improves the computing efficiency. Step III applies reversible jump sampling to transit between 𝔐_0_ and 𝔐_1_. Finally, Step IV updates the time deviance Δ*t*. We use the notation *P*∗(.) to represent the full conditional posterior probability for a given parameter, and define 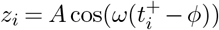 (with *z*_*i*_ = 0 for 𝔐_0_).

#### Step I: Gibbs sampling for *M* and *σ*^2^

The full conditional posterior distributions of *M* and *σ*^2^ can be derived and updated using Gibbs sampling:

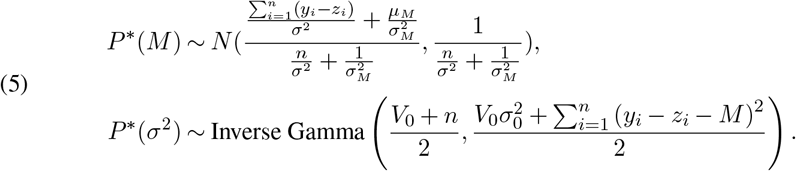

#### Step II: Update within 𝔐_1_: slice sampling for joint update of *A* and *ϕ*

The joint posterior distribution of *A* and *ϕ* is

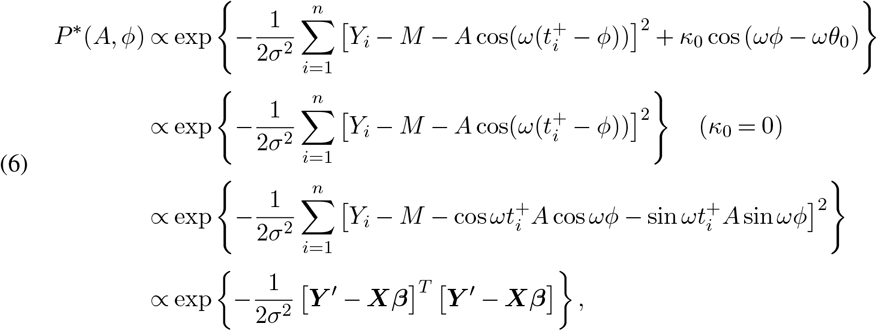

where

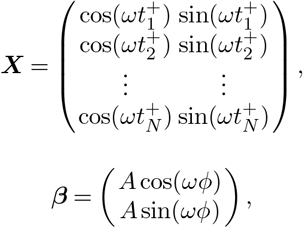

and

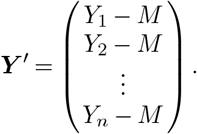

If we re-parameterize the above density as a posterior density for ***β*** with

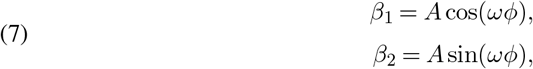

then the inverse function can be obtained as

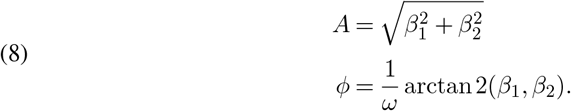

and the corresponding Jacobian determinant can be calculated as

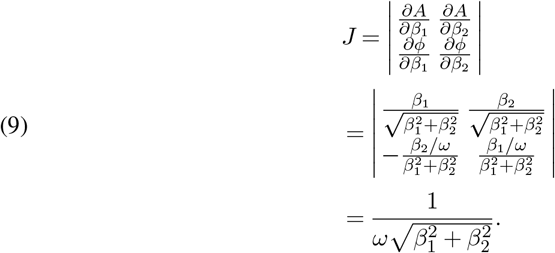

Thus the transformed conditional posterior density of ***β*** is

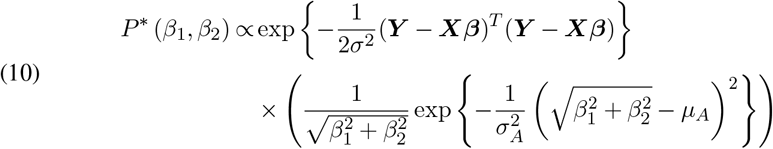

Now we can perform slice sampling as follows:

1. Sample *u* from 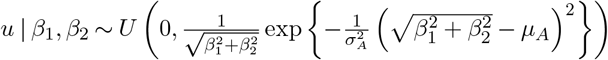
2. Sample (*β*_1_, *β*_2_) from

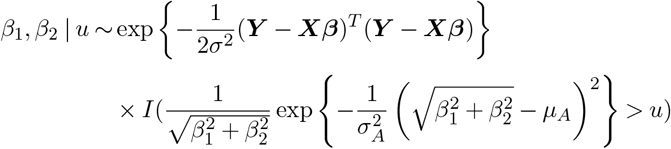

The function

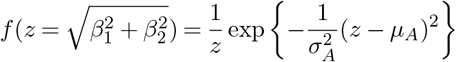

is a decreasing function on *z >* 0, so if we solve for 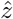 s.t. *f* (*z*) = *u*, any *z* with 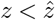 will satisfy the constraint.

Finally, Step II can be achieved by a two-step Gibbs sampling:

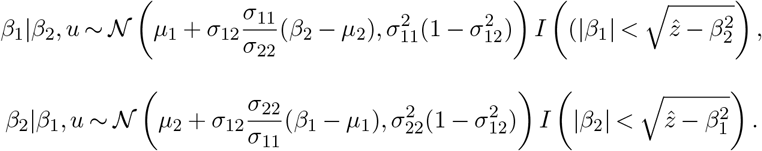

#### Step III: Update between 𝔐_1_ and 𝔐_0_ using reversible jump sampling

Next, we show the between model updating procedure using reversible jump. For reversible jump, when selecting between models with different dimensions, we use augmenting variables to match the parameter space dimension across models. The parameter space of the two models that we want to select from are 𝔐: ***β***_**0**_ **=** {*M* ^′^ *σ* ^2′^} and 𝔐_1_: ***β***_**1**_ ={*M, σ*^2^, *A, ϕ*.}

Suppose the current state is (𝔐 _0_, ***β***_**0**_), and we propose augmenting variables **u** = {*A, ϕ*.} The acceptance ratio of reversible jump is calculated as below:

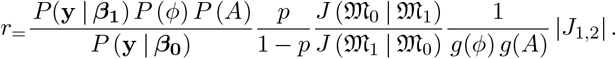

In this formula, *p*, as the prior of *P* (*ρ* = 1), is set to *p* = 0.2 to reflect the prior knowledge of proportion of circadian biomarkers. The probability of transition proposal *J* between models is set to 0.5 in both direction, resulting in 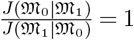. This indicates that the probability of transitioning from model 𝔐_0_ to 𝔐_1_ is equal to the probability of transitioning from model 𝔐_1_ to 𝔐_0_. Furthermore, |*J*_1,2_ |represents the Jacobian of the one-to-one transformation from the parameter space {***β***_**0**_, **u** }under model 𝔐_0_ to the parameter space ***β***_**1**_ under model 𝔐_1_. Since no parameter transformation is performed between the two models, the Jacobian determinant equals 1. Finally, the *g*(*A*) and *g*(*ϕ*) are the proposal probability of the two parameters, which we set to be their prior distritions.

After simplification by plugging the above values, we get

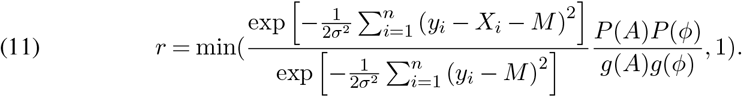

The sampling procedure is as follows, with probability 0.5, the chain stays at the current state and performs corresponding samplings, otherwise, the chain will jump to the other state with the acceptance ratio *r* if we are jumping from 𝔐_0_ to 𝔐_1_ and 1/*r* otherwise.

#### Step IV: Update Δ*t*_*i*_ in BayCT

Finally, the posterior distribution of Δ*t*_*i*_ in the BayCT model is updated as:

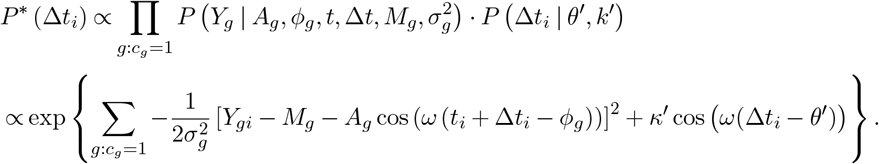

Here, the set of genes with index *c*_*g*_ = 1 is chosen to update Δ*t*. Without prior knowledge, we update *g*: *c*_*g*_ = 1 as the rhythmic genes identified in the last MCMC iteration, i.e.,{*g*: *c*_*g*_ = 1} = {*g*: *ρ*_*g*_ = 1 }. In real application, we pre-select a small set of core clock genes with known strong circadian rhythmicity, denoted by *s*_*g*_ = 1, and Δ*t* is updated with genes that are both pre-selected and estimated as rhythmic, i.e., {*g*: *c*_*g*_ = 1} = {*g*: *s*_*g*_ · *ρ*_*g*_ = 1}.

If we denote *P* ^∗^ (Δ*t*_*i*_) ∝ exp {*f* (Δ*t*_*i*_)}, the sampling procedure is as following:

1. Draw *u* ∼ *U* (0, exp {*f* (Δ*t*_*i*_)}), which is equivalent to firstly drawing *z* ∼ exp(1) and then calculate log *u* = *f* (Δ*t*_*i*_) – *z*
2. Calculate the range of 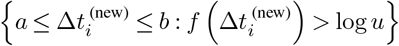
3. Draw 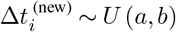

### 2.3. BayCT-M model

The BayCT-M extends BayCT model from single brain region to multiple brain regions. We denote the expression level of gene *g* (*g* = 1, 2, …, *G*) in brain region *j* (*j* = 1, 2, …, *J*) and subject *i* (*i* = 1, 2, …, *n*_*j*_) at time point *t*_*ji*_ as *Y*_*gji*_. In the particular setting of our motivating example, data from the three brain regions are from *N* = 59 identical subjects. As such, *n*_1_ = *n*_2_ = *n*_3_ = *N* = 59 and *t*_1*i*_ = *t*_2*i*_ = *t*_3*i*_ = *t*_*i*_. Notably, our implementation in R also allows for partially overlapped subjects across brain regions, i.e.,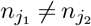 for some *j*_1_, *j*_2_. The cosinor model is parameterized as:

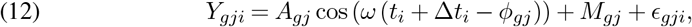

where 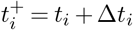 is the MCT. The priors are the same as in BayCT, except that *ϕ*_*gj*_ shares a common mean *θ*_*g*_ in the von Mises prior across brain regions. The *A*_*gj*_, 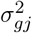, and *M*_*gj*_ have more considerable variance across groups and thus are assigned independent priors for each tissue (Zhang et al., 2014; Mure et al., 2018; Talamanca, Gobet and Naef, 2023):

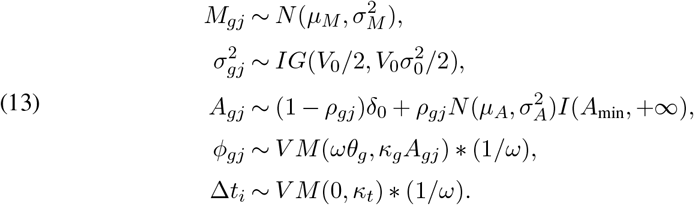

Similar to BayCT, the indicator variable *ρ*_*gj*_ = 1 indicates gene *g* in brain region *j* is rhythmic and *ρ*_*gj*_ = 0 otherwise. We set 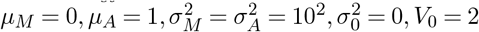 and *κ*_*t*_ = 6.The dependence of *ϕ*_*gj*_ prior on *A*_*gj*_ is grounded in the rationale that a smaller *A*_*gj*_ implies a weaker signal and thus a greater dispersion in *ϕ*_*gj*_. We further define the conjugate joint hyperprior for the hyperparameters *θ*_*g*_ and *κ*_*g*_:

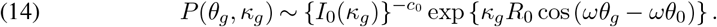

In this prior distribution, *I*_0_(*κ*_*g*_) is the Bessel function of the first kind with order 0. The hyperprior indicates *c*_0_ number of prior observations in the direction *θ*_0_, and *R*_0_ corresponds to the component projected along *θ*_0_ (i.e., in the known direction) of the resultant of *c*_0_ observations (Damien and Walker, 1999). We set *c*_0_ = 1, *R*_0_ = 1, and *θ*_0_ = 0, which constitutes a weakly informative prior, providing a baseline orientation while allowing the data to largely influence the posterior estimates. The corresponding Gibbs sampling procedure proposed in (Damien and Walker, 1999) is used for updating *θ*_*g*_ and *κ*_*g*_. Figure 1B shows the graphical representation of BayCT-M under 𝔐_0_(*ρ*_*gj*_ = 0) and 𝔐_1_(*ρ*_*gj*_ = 0). The whole MCMC sampling procedures for the BayCT-M are detailed in Supplement Section 1 Posterior Inference. Inference outputs (posterior mean and standard deviation) of four parameters from BayCT-M are: (1) *ρ*_*gj*_: circadian indicator for gene *g* in tissue *j*, (2) *θ*_*g*_: canonical phase of gene *g*, (3) *ϕ*_*gj*_: tissue-specific phase for a circadian gene *g*, (4) Δ*t*_*i*_: deviation of molecular circadian time of subject *i*.

## 3. Simulations

### 3.1. Simulation to evaluate BayCT model

We generate data for 1000 genes, consisting of 200 rhythmic and 800 arrhythmic genes, to reflect the higher prevalence of arrhythmic genes in real applications. For the rhythmic genes, we set amplitude *A* = 0.6, 0.8, 1, with a constant variance *σ*^2^ = 1. The time deviation is varied along two dimensions: the magnitude of deviation |Δ*t* |, set at 0, 1, 2, and 4 hours, and the proportion of the total 30 samples affected by this deviation, set at *r* = 0, 20%, 40%, with equal numbers of samples having positive and negative |Δ*t*| values. Each setting is repeated 30 times.

We first evaluate MCT estimation accuracy, comparing BayCT with various methods, including TimeSignature (TS), ZeitZeiger (ZZ), CHIRAL, CYCLOPS, and Cyclum. Note that CHIRAL, CYCLOPS and Cyclum are not standard supervised machine learning methods. These tools are unsupervised without utilizing the input TOD. They, however, can generate pseudo-time, which is prediction of the MCT of each subject with certain unknown shift. Consequently, these three methods are not applicable to perform cross-validation in the MCT prediction evaluation. In Figure 2, the prediction accuracy is calculated by using all subjects to construct the model and then applied to the same subjects. This may cause slight overfitting in the accuracy evaluation but it gives a fair method comparison. To avoid overfitting, Figure S1 conducts five-fold cross validation and compare methods excluding CHIRAL, CYCLOPS and Cyclum.

**Fig 2.**
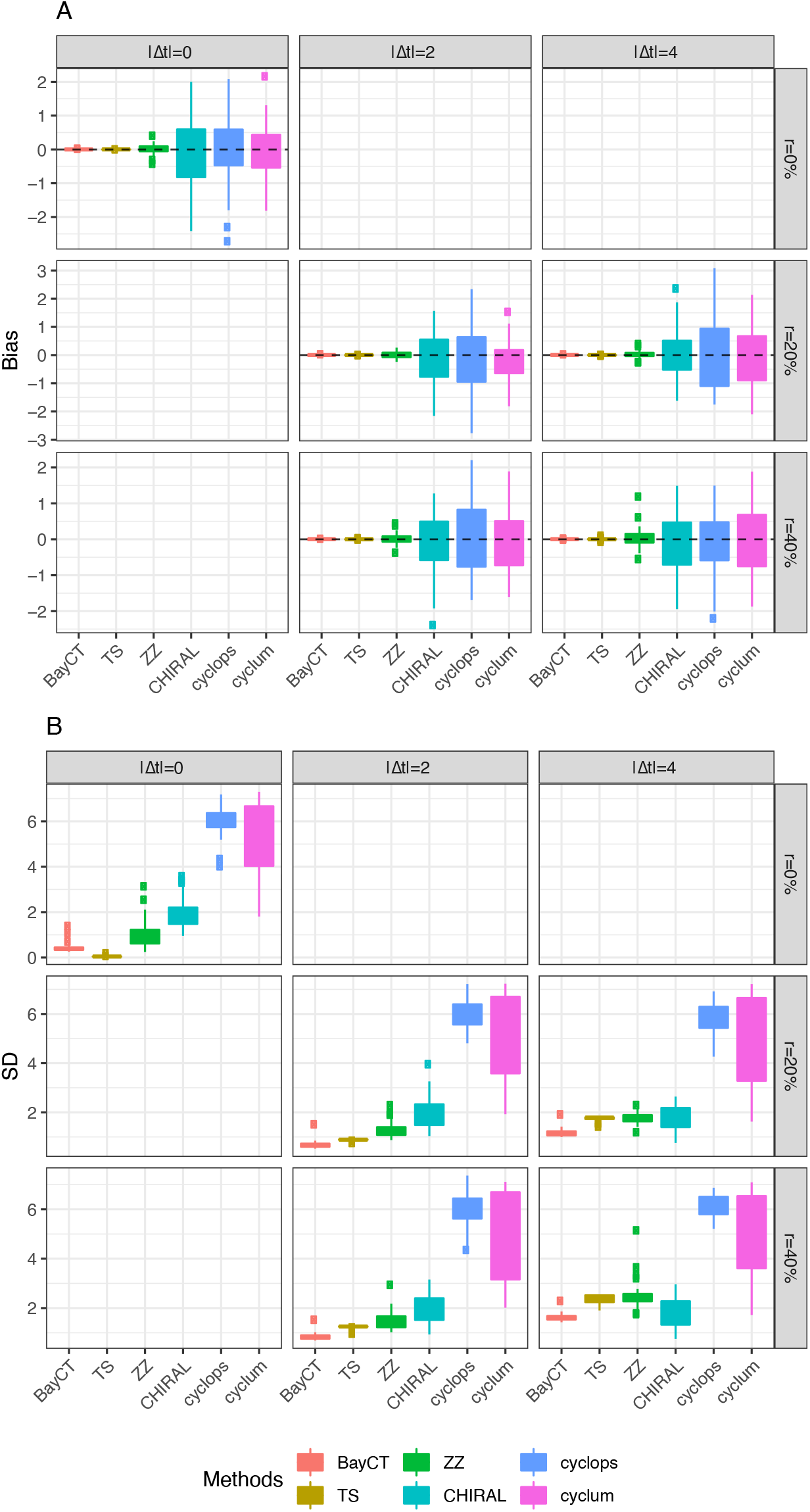
Bias and standard deviation (SD) of the MCT estimations.

Next, we compare biomarker detection accuracy among BayCT, BayC and the Cosinor model, using the area under the Receiver Operating Characteristic curve (AUC) as a threshold-free measure. For Bayesian methods, sensitivity and specificity are calculated with varying cutoffs for *P* ^∗^(*ρ* = 1), while the frequentist Cosinor method uses varying p-value cutoffs. Additionally, we assess the parameter estimation consistency through the bias and standard deviation of the point estimators from each method.

#### MCT estimation accuracy

The simulations in Figure 2A shows that BayCT and TS give the most accurate MCT prediction, regardless of Δ*t* and *r*. In Figure 2B, when there is no time deviation (i.e., Δ*t*_*i*_ = 0 for all *i*), BayCT has slightly larger standard deviation (SD) than TS due to slight loss of efficiency by estimating Δ*t*. But when time deviation indeed exist, BayCT outperforms TS as well as other methods. Larger and more prevalent time deviations (e.g., Δ*t* = 4) lead to more significant superiority in BayCT.

An extra advantage of BayCT over other machine learning methods is its inference of a Bayesian credible interval for the MCT prediction uncertainty. Supplement Figure S2 shows the actual coverage probability when the 95% credible interval of MCT contains the truth. The result shows general accurate estimation of the 95% credible intervals.

#### Circadian biomarker detection

Figure 3A presents the area under the curve (AUC) evaluation of circadian gene detection using BayC, BayCT and frequentist Cosinor model. In the baseline scenario where Δ*t*_*i*_ = 0 for all *i* (i.e., when MCT aligns perfectly with TOD for all samples), BayC and Cosinor achieve comparable high accuracy. The result is consistent with the general observation that Bayesian approaches and their frequentist counterparts often achieve similar high performance when both modelled adequately. BayCT has a slightly lower AUC performance, due to the slight loss of efficiency from estimating Δ*t*_*i*_ when they are all zero. However, as the degree of deviance grows, BayCT becomes increasingly effective, surpassing the other two methods. For example, when *A* = 0.8, and *r* = 40% of samples have |Δ*t*| = 4, BayeCT achieves mean AUC=0.909, compared to mean AUC of 0.869 and 0.852 for BayC and Cosinor. Notably, AUC of BayCT remains almost at the same level for fixed *A*, showing effective time deviation correction and the robustness provided by BayCT.

**Fig 3.**
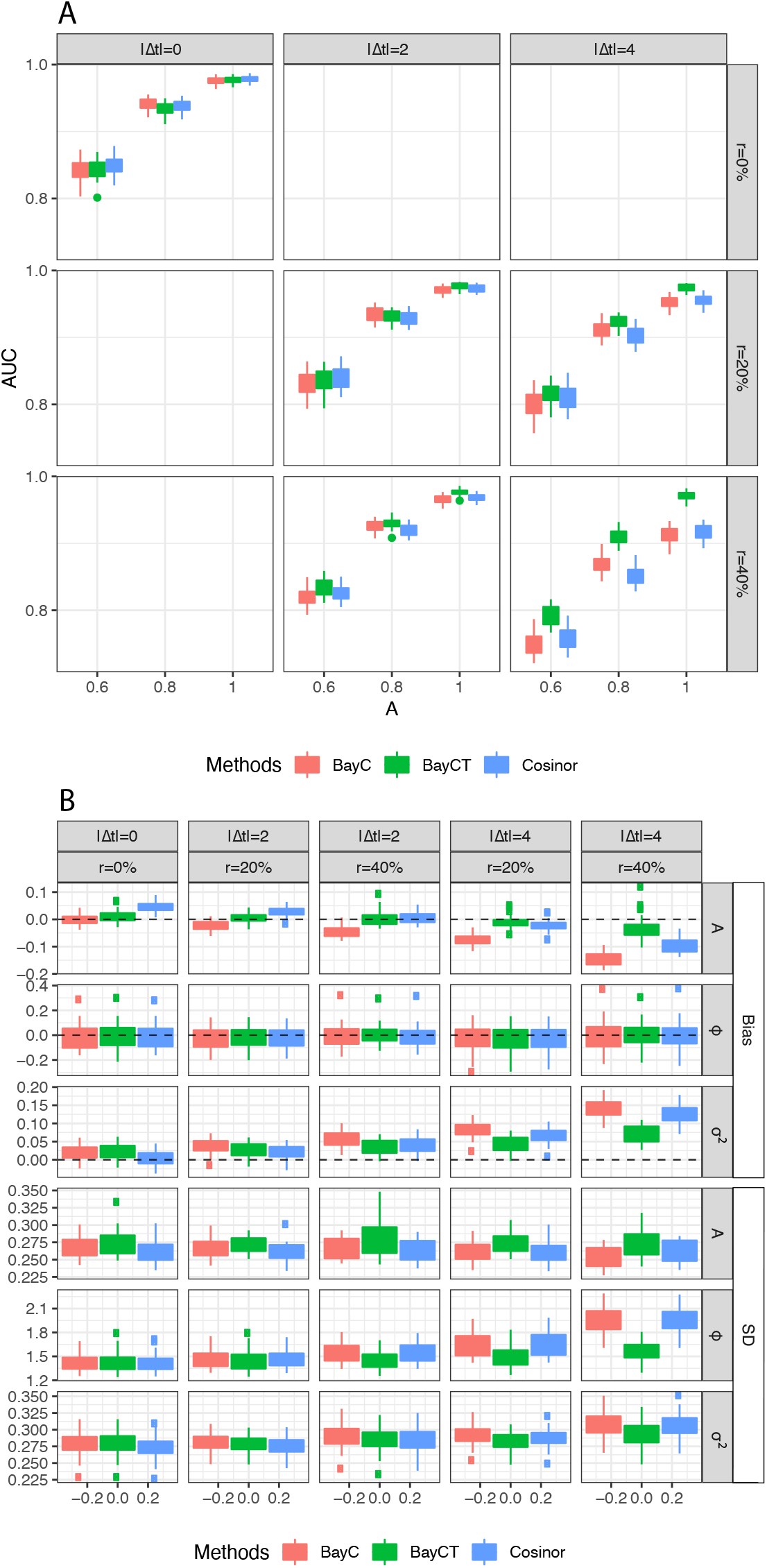
A. Comparison of rhythmicity accuracy of the three methods with AUC. The columns, moving from the left to right, represent an increasing magnitude of Δt, while the rows, from top to bottom, present an increasing percentage of samples affected by Δt. B. Bias and SD evaluation for the three methods when simulating A = 0.8.1

We further assess estimation accuracy in terms of bias and standard deviation of amplitude A, phase *ϕ*, and expression error term *σ*^2^ for BayC, BayCT and Cosinor. The result at simulating *A* = 0.8 is presented in Figure 3B. In terms of estimating A and *σ*^2^, BayCT maintains an unbiased estimation performance, whereas both BayC and Cosinor tend to underestimate A and overestimate *σ*^2^ with increasing proportion of samples with time deviation, *r*. Since *A*/*σ* stands for signal-to-noise ratio, this result can be explained by that time deviation introduces uncertainty in signal detection. In terms of *ϕ*, all three methods are unbiased when *r* increases. BayC and Cosinor, however, have much larger SD compared to BayCT when *r* increases.

### 3.2. Simulations to evaluate BayCT-M model

We simulate 2000 genes in 3 brain regions from 30 samples, that is, *G* = 2000, *J* = 3, and *n*_*j*_ = 30. Then, we simulate data for 1300 arrhythmic genes in all 3 regions, 100 genes rhythmic only in each tissue (300 genes in totla), 100 genes rhythmic in any two tissues (300 genes in total), and 100 genes rhythmic inall 3 regions. We set amplitude *A*_*gj*_ = 0.6, 0.8, 1 and a constant variance 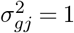 to generate adequate intrinsic effect size (i.e., *A/σ*) for sufficient statistical power (Zong et al., 2023). Similar to the previous subsection, the circadian time deviation varies in two dimensions: 1) the magnitude of deviation, |Δ*t*| ∈ 0, 2, 4. 2) the proportion of total samples affected by this deviation was set at *r* = 0%, 20%, 40%, with equal samples with positive and negative |Δ*t*| values. We categorize simulations into two settings: 1) SamePhase: *ϕ*_*gj*_ are same across tissues for rhythmic genes. 2) DiffPhase: *ϕ*_*gj*_ are different across tissues for rhythmic genes with magnitude 2 and 4, i.e., *ϕ*_*g*2_ −*ϕ*_*g*1_ = 2 and *ϕ*_*g*3_ −*ϕ*_*g*1_ = 4. Each combination of these variations is simulated 30 times. The performance of BayCT-M in the two settings are highly similar, demonstrating robust capacity of BayCT-M under heterogeneity.

#### MCT estimation accuracy

Supplement Figure S3 and Figure S5 show bias and SD of MCT estimates under SamePhase and DiffPhase simulations, respectively. BayCT-M consistently gives unbiased estimates regardless of |Δ*t*| and *r*. As amplitude *A* increases, BayCT-M gives more accurate estimates with smaller SD, indicating that the model is efficient if we have strong rhythmicity signal.

To assess the efficiency of BayCT-M relative to BayCT in estimating Δ*t*, we apply BayCT separately on three tissues from the simulated data and compute the SD ratio of BayCT-M to BayCT, as shown in Figure 4. Notably, the efficiency gain is most pronounced at lower amplitude (*A* = 0.6), where the SD ratio drops below 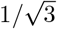 (i.e., the expected asymptotic optimal efficiency). As the amplitude increases, these ratios converge to 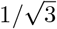. The result shows that BayCT-M is asymptotically efficient in integrating information across brain regions for predicting MCT.

**Fig 4.**
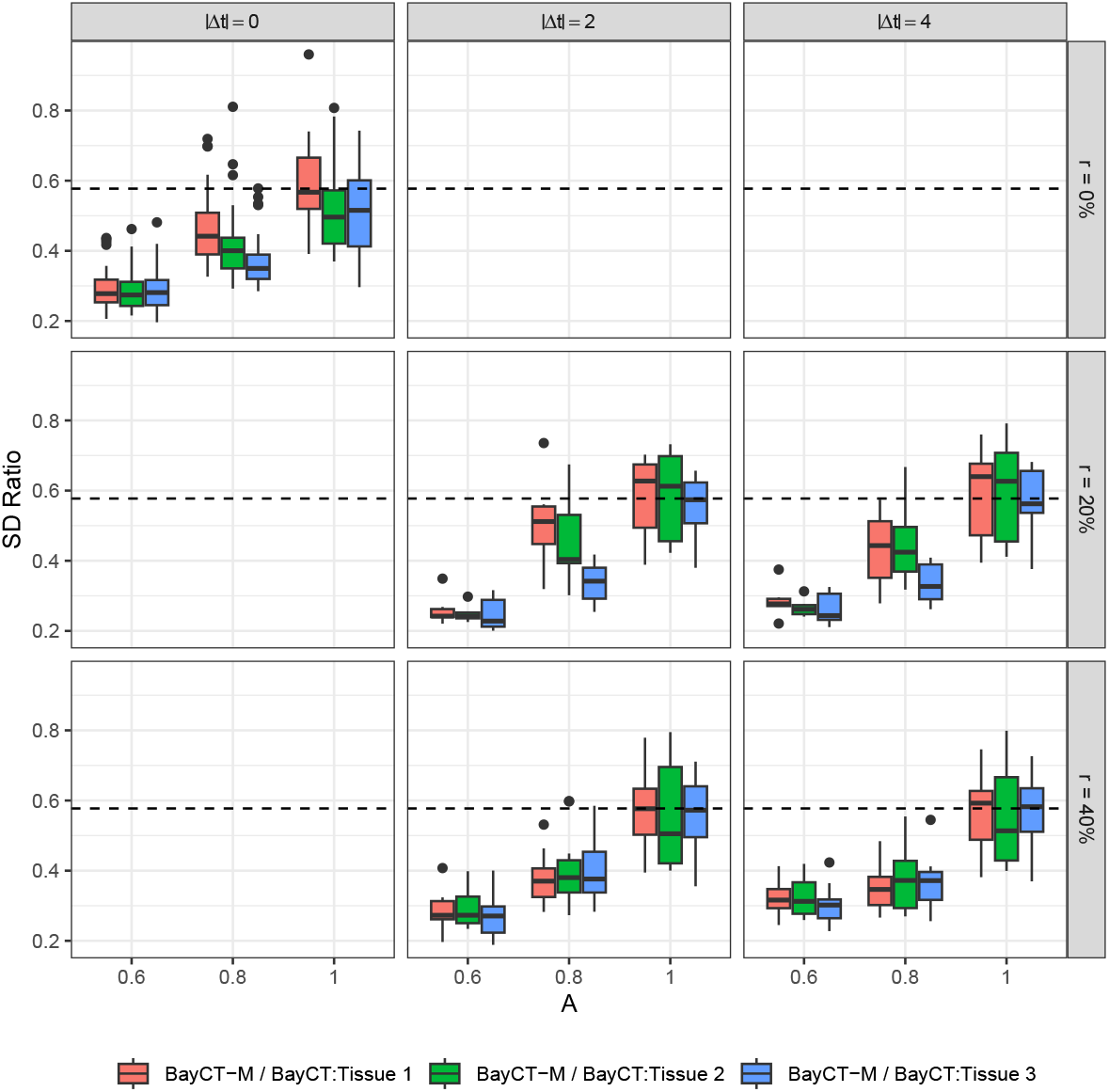
The SD ratio of BayC T-M to BayCT on each tissue with increasing amplitude A when estimating Δt. The horizontal dashed line is 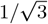 The columns, moving from the left to right, represent an increasing magnitude of Δt, while the rows, from top to bottom, present an increasing percentage of samples affected by Δt.

#### Circadian biomarker detection

The evaluation on circadian gene detection accuracy in Figure S4 and Figure S6 show that BayCT-M is capable to identify circadian gene with high performance. The AUC approaches 1 when the signal is strong enough (i.e., *A* = 1). Comparisons in Figure S7 confirm that BayCT-M performs comparably to the single-tissue BayCT method. BayCT-M shows a slight advantage at lower signal amplitudes, indicating that integrating multi-tissue information preserves robust biomarker detection.

## 4. Real applications

In this section, we evaluate performance of BayCT and BayCT-M with two real datasets. In the first application, since biological/genetic and environmental variations among mice are minimal, we assume TOD=MCT (i.e., Δ*t* = 0) in the mouse study. We simulate Δ*t* to generate semi-synthetic data from real data to evaluate MCT prediction performance of BayCT and BayCT-M. In the second application, we apply BayCT to a human postmortem transcriptomic dataset with three brain regions for both biomarker detection and MCT estimation.

### 4.1. Evaluation in mouse data with synthetic Δt

Circadian gene markers have been found to be tissue-specific. As such, a transcriptomic study in mouse by Zhang et al. (2014) provided an circadian atlas composing 12 tissues, each sampled in every 2 hours for 2 days (24 samples per tissue). Since all mice were entrained with the same schedule before the sample collection, the MCT is considered to be aligned perfectly to the experiment time (i.e., Δ*t* = 0).

After filtering out probes with negative values and duplication, 35,556 genes (transcripts) are left and we further normalize the data with quantile normalization. To assess the MCT prediction methods on real data, we generate pseudo datasets by artificially introducing noise to MCT. Specifically, for a randomly selected half of the samples, we add Δ*t* to the observed time, while for the other half, we subtract Δ*t* from the observed time to create the new observed TOD. This process simulates scenarios where the recorded TOD might deviate from the underlying MCT due to various factors.

Then we apply the BayCT and other MCT prediction methods to the data using the pseudo time as the input. For BayCT, we update |Δ*t*| for the 12 tissues separately with the 11 present core rhythmic genes: NPAS2, PER2, CRY1, CIART, PER1, NR1D1, NR1D2, TEF, CRY2, PER3 and DBP, while simultaneously estimating the rhythmicity parameters for all the genes. The results show that the 95% credible intervals (CIs) covered 94.4% of the artificial Δ*t*, indicating the model’s ability to accurately capture the introduced noise. Additionally, we apply other MCT prediction methods with the same set of 11 core clock genes, using procedures described in Section 3. As shown in Figure 5A, the comparison yielded results similar to those from our simulation, with BayCT model exhibiting consistent unbiasedness and small variance overall. These findings demonstrate the BayCT’s robust performance on real data.

**Fig 5.**
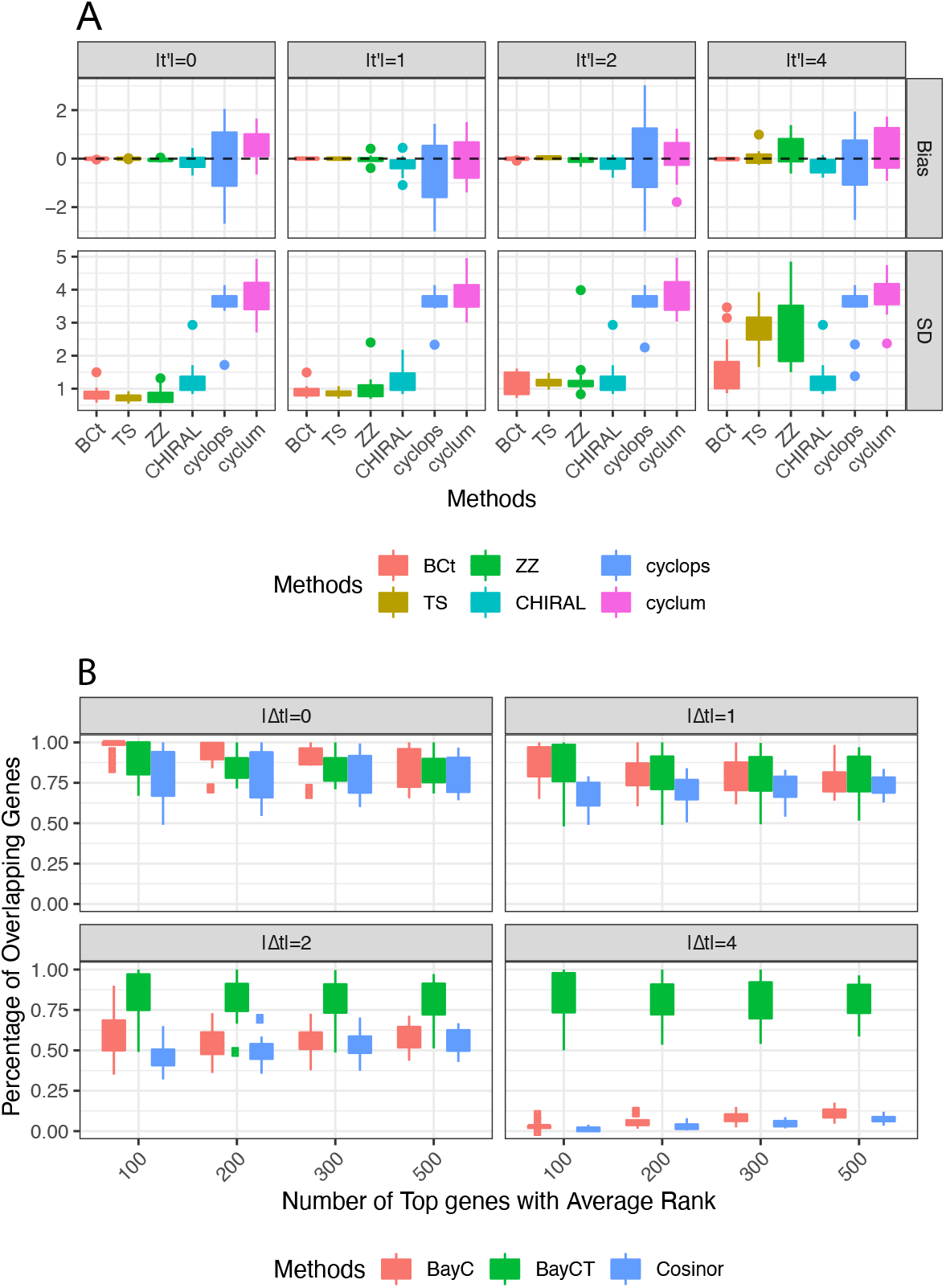
A. Bias and SD evaluation for MCT prediction, pooled over 12 tissues. B. The overlapping of top rhythmic genes for BayC, BayCT, and Cosinor with increasing artificial Δt.

Subsequently, we evaluate the accuracy of rhythmicity identification of BayC, BayCT, and Cosinor with pseudo |Δ*t*| = 1, 2, 4. To obtain a reference rank of gene rhythmicity which we can compare to, we applied the three models on the original data (when |Δ*t* = 0|) and obtain the gene rhythmicity rank for each method. For BayC and BayCT, genes with larger *P* ^∗^(*ρ* = 1) have smaller ranks, while for Cosinor, genes with smaller p-value have smaller ranks. The “ensemble reference” top genes were ordered by the smallest average rank across the three methods. We then examin the percentage of the n-top-ranked rhythmic genes from the “ensemble reference list” that are overlapping with the n-top-ranked rhythmic genes from each method, with *n* = 100, 200, 300, and 500. Figure 5B shows that when |Δ*t*| = 0, BayC has the highest overlapping percentage. However, as |Δ*t*| increases, both BayC and Cosinor lost accuracy in identifying the top rhythmic genes. In contrast, BayCT displays robust performance across varying |Δ*t*|, with the results being comparable to results at |Δ*t*| = 0. This demonstrates the BayCT’s overall accurate and robust identification of circadian genes under the impact of varying |Δ*t*|.

### 4.2. Human postmorten data with three brain regions

In the second case study, we apply BayCT and BayCT-M to RNA-seq data from Ketchesin et al. (2021), which involves samples from three distinct brain regions - nucleus accumbens (NAc), caudate and putamen - of the same 59 subjects.

We independently apply BayCT to each of the three brain regions and use BayCT-M on all three brain regions. Figure 6A shows the BayCT-M Δ*t* ordered by the mean with the corresponding 90% credible interval. All of the BayCT-M Δ*t* have a 90% credible interval width of less than 2.4 hours, showing high concordance of Δ*t* estimates across three brain regions.

**Fig 6.**
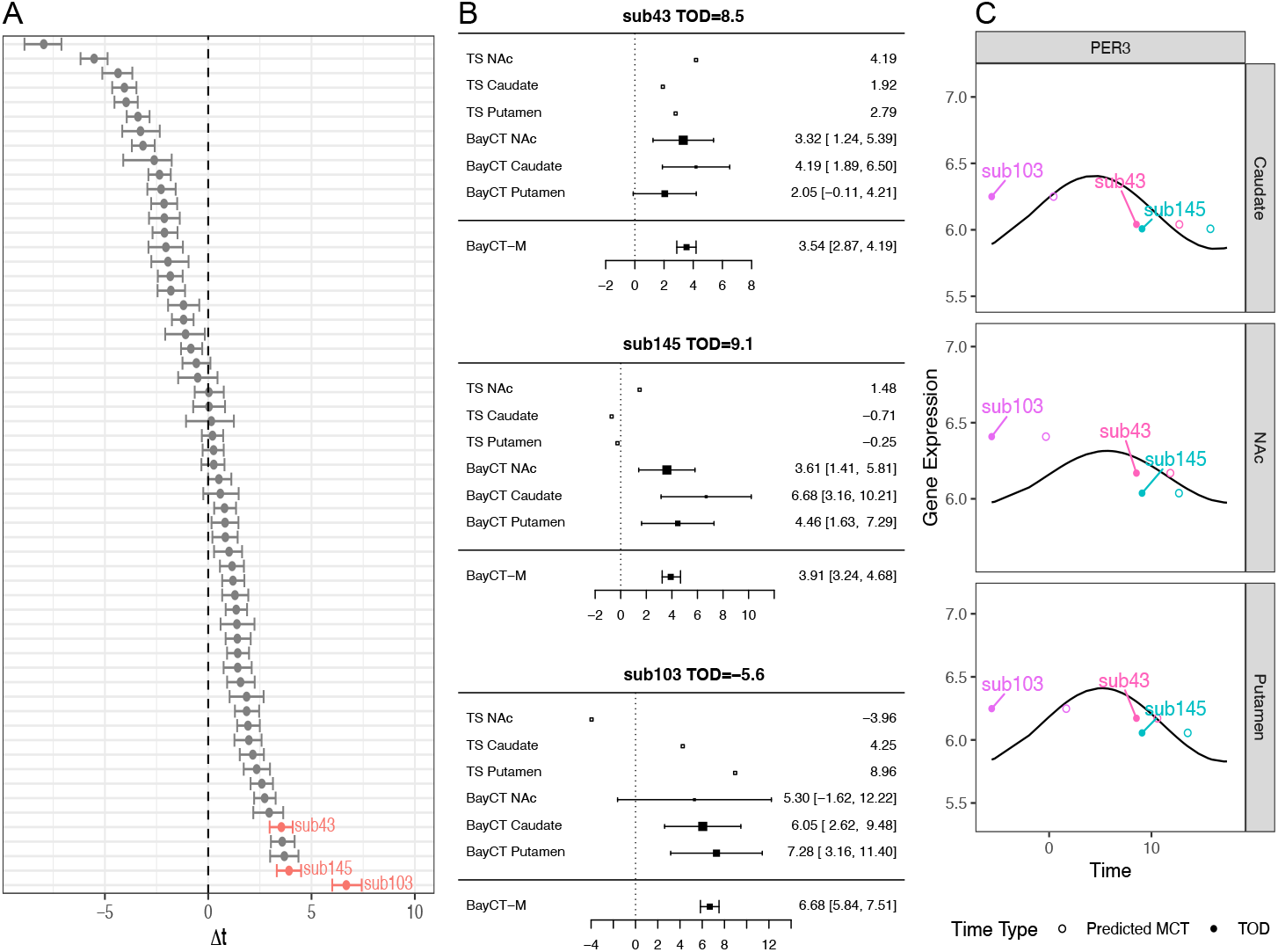
The BayCT, BayCT-M, and TS Δt estimates for three brain regions. A. BayCT-M Δt estimates and 90% credible interval for all subjects. B. Forest plots of subject 43, 145, and 103. C. Circadian gene expression pattern of PER3 in Caudate, Nac, and Putamen. x axis is TOD or predicted MCT and y axis is gene expression level. The black line is the fitted sinusoidal curve using TOD

For further comparison, we compare these BayCT-M Δ*t* estimates with those obtained from TimeSignature (TS), which is identified as the second-best performer in the extensive simulation. Remarkably, the comparison reveals a high degree of similarity, with 54 out of 59 (91.5%) samples exhibiting a difference of less than 44 hours (Figure S8). For demonstration, Figure 6B provides forest plots of BayCT Δ*t* in three subjects, BayCT-M Δ*t*, and the Δ*t* estimations from TS. Since TS is a machine learning method, only the Δ*t* point prediction is available. In contrast, BayCT and BayCT-M provides uncertainty quantification by 95% credible interval, in addition to the mean point estimation. In Subject 43, the mean Δ*t* = 3.54 and narrow Δ*t* credible interval = [2.87, 4.19] from BayCT-M shows high inference confidence of the difference. TS predictions in three brain regions gives consistent conclusions of Δ*t* = 4.19, 1.92 and 2.79, respectively. In subject 145, BayCT-M also concludes highly confident Δ*t* with mean = 3.91 and credible interval = [3.24, 4.68]. In contrast, TS shows inconclusive Δ*t* prediction at 1.48, −0.71 and −0.25. In Subject 103, BayCT concludes consistent signals for Δ*t* across three brain regions although NAc has larger variance. As such, the BayCT-M result gives significant conclusion of mean Δ*t* = 6.68 and credible interval = [5.84, 7.51]. Prediction result from TS is relatively consistent with BayCT in Caudate and Putamen (Δ*t* = 4.25 and 8.96) while very different in NAc (Δ*t* = −3.96). Again TS can only provide predication without uncertainty quantification in the prediction compared to BayCT. We also notice that BayCT-M gives us much narrower credible interval in this real application. This is possibly due to finite sample or data noise and our simulation results in Figure 4 guarantees that BayCT-M Δ*t* estimate has a reasonable efficiency when rhythmicity signal is strong. In Figure 6C, we use a core clock gene PER3 to illustrate how MCT adjusts from TOD to provide better circadian curve fitness. PER3 appears to be a circadian genes in all three brain regions (posterior probability of *ρ*_*gj*_ = 1 is 1, for *j* = 1, 2, 3). Expression levels plotted with MCT (circles) fit the circadian curves much better than TOD (solid dots), except for poor fitness in both MCT and TOD for NAc in Subject 103. This raw data visualization confirms the likely time deviations Δ*t* in these three subjects.

## 5. Discussion

The inherent biological variability of MCT presents significant challenges to not only the chronotherapy but also the broader researches in human circadian biology. Traditional studies often equates the TOD with MCT, introducing measurement noise into the analyses. On the other hand, existing time prediction methods focus on estimating MCT, but not on inferences of circadian biomarkers and their associated rhythmic parameters.

In response to these challenges, we developed BayCT, a novel Bayesian circadian detection model that precisely adjusts the discrepancies between the observed time and MCT. The model employs full Bayesian inference, enabling the inference of both the rhythmic parameters in detected biomarkers and the uncertainty in MCT estimation. BayCT has shown superior performance in circadian biomarker detection compared to the traditional frequentis method Cosionor and BayC without MCT adjustment. It also provides more accurate MCT estimation than other machine learning approaches. Moreover, BayCT’s extension to BayCT-M enables us to deal with multi-tissue data within a full Bayesian framwork.

The application of BayCT to real datasets further validates its effectiveness. By introducing pseudo Δ*t* into the mouse data, we are encouraged by its robust performance in estimating the MCT and rhythmicity identification across various levels of noise strength. Additionally, the concordant Δ*t* estimations across multiple human brain tissues demonstrates the reliability of its results.

In our motivating example, data from the three brain regions are from *N* = 59 identical subjects. However, in our R package, BayCT, the BayCT-M method can implement the general situation when data across brain regions or organs are from partially overlapped subjects.

BayCT is designed assuming Gaussian distributed residual errors, making it applicable for a variety of omics data types, such as transcriptomics, proteomics, and log-transformed methylation data. For the human postmorten RNA-seq data from Ketchesin et al. (2021) with *G* = 14, 540 transcripts, *J* = 3 brain regions, and *n*_*j*_ = 59 subjects, BayCT-M takes 4.45 hours to complete 3,000 MCMC simulations on a personal laptop with 18GB RAM. The computing is expected to significantly improve if calling Rcpp from R.

In summary, BayCT enhances circadian analysis by adjusting for discrepancies between the MCT and observed TOD via a full Bayesian inference framework. It presents robust circadian detection and inference with varying levels of biological variation of MCT, which enhance the power and accuracy of circadian analysis.

## Acknowledgments

The authors would like to thank the anonymous referees, an Associate Editor and the Editor for their constructive comments that improved the quality of this paper.

## SUPPLEMENTARY MATERIAL

### Supplement to

The supplement contains detailed posterior inference procedures for the BayCT and BayCT-M models, along with additional simulation results and supplementary figures.

