## Supplementary Material for "Bayesian rhythmic model for jointly detecting circadian biomarkers and predicting molecular circadian time in human post-mortem brain transcriptome"

**1. Posterior Inference.** This section presents details for some steps in our sampling procedures in BayCT and BayCT-M model.

1.1. *BayCT model.* For computational simplicity, we re-parameterize the full conditional $P^*(\Delta t_i)$ in **Step IV: Update $\Delta t_i$ in BayCT** as the following:

(1)

$$P^*(\Delta t_i) \propto \exp\left\{\sum_{\rho_g=1} -\frac{1}{2\sigma_g^2}\left[Y_{gi} - M_g - A_g\cos\left(\omega\left(t_i - \phi_g + \Delta t_i\right)\right]^2 + \kappa'\cos\left(\omega\left(\Delta t_i - \theta'\right)\right)\right\}$$

$$\propto \exp\left\{\sum_{\rho_g=1} -\frac{1}{2\sigma_g^2}\left[Y_{gi} - M_g - X_{gi1}\cos\omega\Delta t_i - X_{gi2}\sin\omega\Delta t_i\right]^2 + \kappa'\cos\omega\theta'\cos\omega\Delta t_i + \kappa'\sin\omega\theta'\sin\omega\Delta t_i\right\}$$

$$\propto \exp\left\{-\frac{1}{2}\sum\frac{X_{gi1}^2}{\sigma_g^2}\cos^2\omega\Delta t_i - \frac{1}{2}\sum\frac{X_{gi2}^2}{\sigma_g^2}\sin^2\omega\Delta t_i - \sum\frac{X_{gi1}X_{gi2}}{\sigma_g^2}\sin\omega\Delta t_i\cos\omega\Delta t_i\right.$$

$$\left. + \left(\sum\frac{\left(Y_{gi} - M_g\right)X_{gi1}}{\sigma_g^2} + \kappa'\cos\omega\theta'\right)\cos\omega\Delta t_i + \left(\sum\frac{\left(Y_{gi} - M_g\right)X_{gi2}}{\sigma_g^2} + \kappa'\sin\omega\theta'\right)\sin\omega\Delta t_i\right\}$$

$$\propto \exp\left\{R_1\cos\left(\omega\Delta t_i - \alpha_1\right)\right\}\exp\left\{R_2\cos\left(2\omega\Delta t_i - \alpha_2\right)\right\},$$

where

[*]George C. Tseng is corresponding author

$$X_{gi1} = A_g \cos(\omega(t_i - \phi_g)),$$
$$X_{gi2} = A_g \sin(\omega(t_i - \phi_g)),$$
$$R_{1i}^2 = \left(\sum \frac{(Y_{gi} - M_g)\, X_{gi1}}{\sigma_g^2} + \kappa' \cos\omega\theta'\right)^2 + \left(\sum \frac{(Y_{gi} - M_g)\, X_{gi2}}{\sigma_g^2} + \kappa' \sin\omega\theta'\right)^2,$$
$$\tan\alpha_{1i} = \frac{\sum \frac{(Y_{gi}-M_g)X_{gi2}}{\sigma_g^g} + \kappa' \sin\omega\theta'}{\sum \frac{(Y_{gi}-M_g)X_{gi1}}{\sigma_g^2} + \kappa' \cos\omega\theta'},$$
$$R_{2i}^2 = \left(-\frac{1}{4}\sum \frac{X_{gi1}^2}{\sigma_g^2} + \frac{1}{4}\sum \frac{X_{gi2}^2}{\sigma_g^2}\right)^2 + \left(\frac{1}{2}\sum \frac{X_{gi1}X_{gi2}}{\sigma_g^2}\right)^2,$$

and

$$\tan\alpha_{2i} = \frac{-\frac{1}{2}\sum \frac{X_{gi1}X_{gi2}}{\sigma_g^2}}{-\frac{1}{4}\sum \frac{X_{gi1}^2}{\sigma_g^2} + \frac{1}{4}\sum \frac{X_{gi2}^2}{\sigma_g^2}}.$$

Then, we have $P^*(\Delta t_i) \propto \exp\{R_1 \cos(\omega\Delta t_i - \alpha_1) + R_2 \cos(2\omega\Delta t_i - \alpha_2)\}$. And we denote it as $P^*(\Delta t_i) \propto \exp\{g(\Delta t_i)\}$. The sampling procedure is as following:

1. Draw $u \sim U(0, \exp\{g(\Delta t_i)\})$, which is equivalent as first drawing $z \sim \exp(1)$, then calculate $\log u = g(\Delta t_i) - z$.
2. Calculate the range of $\left\{a \le \Delta t_i^{(\text{new})} \le b : g\left(\Delta t_i^{(\text{new})}\right) > \log u\right\}$.
3. Draw $\Delta t_i^{(\text{new})} \sim U(a, b)$

1.2. *BayCT-M model.* In this section, we present the sampling procedure of BayCT-M. The sampling procedures is divided into four steps: Step I occurs only when updating within model $\mathfrak{M}_0$, while Steps I, II, and IV are applied to model $\mathfrak{M}_1$. Step III is performed after each update within either model. Step I updates the two parameters $M_{gj}$ and $\sigma_{gj}^2$ shared by $\mathfrak{M}_0$ and $\mathfrak{M}_1$. Step II uses slice sampling for joint updating of $A$ and $\phi$ in $\mathfrak{M}_1$. Step III applies reversible jump sampling to transit between $\mathfrak{M}_0$ and $\mathfrak{M}_1$. Finally, Step IV updates the time deviance $\Delta t$. We use the notation $P^*(.)$ to represent the full conditional for a parameter, and define $z_{gji} = A_{gj}\cos(\omega(t_i^+ - \phi_{gj}))$ (with $z_i = 0$ for $\mathfrak{M}_0$).

**Step I: Gibbs sampling for $M_{gj}$ and $\sigma_{gj}^2$**

The full conditional posterior distributions of $M$ and $\sigma^2$ can be derived and updated using Gibbs sampling:

$$\begin{aligned} P^*(M_{gj}) &\sim N\left(\frac{\frac{\sum_{i=1}^n (y_{gji} - z_{gji})}{\sigma_{gj}^2} + \frac{\mu_M}{\sigma_M^2}}{\frac{n_j}{\sigma_{gj}^2} + \frac{1}{\sigma_M^2}}, \frac{1}{\frac{n_j}{\sigma_{gj}^2} + \frac{1}{\sigma_M^2}}\right), \\ P^*(\sigma_{gj}^2) &\sim \text{Inverse Gamma}\left(\frac{V_0 + n_j}{2}, \frac{V_0\sigma_0^2 + \sum_{i=1}^n (y_{gji} - z_{gji} - M_{gj})^2}{2}\right). \end{aligned} \tag{2}$$

**Step II: Update within $\mathfrak{M}_1$: slice sampling for $A_{gj}$ and $\phi_{gj}$**

(3)

$$P^*(A_{gj},\phi_{gj}) \propto \exp\left\{-\frac{1}{2\sigma_{gj}^2}\sum_{i=1}^{n_j}\left[Y_{gji}-M_{gj}-A_{gj}\cos\omega t_i^+\cos\omega\phi_{gj}-A_{gj}\sin\omega t_i^+\sin\omega\phi_{gj}\right]^2\right.$$
$$\left.+\kappa_g A_{gj}\cos\left(\omega\phi_{gj}-\omega\theta_g\right)-\frac{1}{2\sigma_A^2}\left(A_{gj}-\mu_A\right)^2\right\}$$
$$\propto \exp\left\{-\frac{1}{2\sigma_{gj}^2}\sum_{i=1}^{n_j}\left[Y_{gji}-M_{gj}-A_{gj}\cos\omega t_i^+\cos\omega\phi_{gj}-A_{gj}\sin\omega t_i^+\sin\omega\phi_{gj}\right]^2\right.$$
$$\left.+\kappa_g A_{gj}\cos\omega\theta_g\cos\omega\phi_{gj}+\kappa_g A_{gj}\sin\omega\theta_g\sin\omega\phi_{gj}-\frac{1}{2\sigma_A^2}\left(A_{gj}-\mu_A\right)^2\right\}$$

where

$$\boldsymbol{X}_j=\begin{pmatrix}\cos(\omega t_1^+) & \sin(\omega t_1^+)\\ \cos(\omega t_2^+) & \sin(\omega t_2^+)\\ \vdots & \vdots\\ \cos(\omega t_{n_j}^+) & \sin(\omega t_{n_j}^+)\end{pmatrix},$$

$$\boldsymbol{\beta}_{gj}=\begin{pmatrix}A_{gj}\cos(\omega\phi_{gj})\\ A_{gj}\sin(\omega\phi_{gj})\end{pmatrix},$$

$$\boldsymbol{Y'}=\begin{pmatrix}Y_{gj1}-M_{gj}\\ Y_{gj2}-M_{gj}\\ \vdots\\ Y_{gjn_j}-M_{gj}\end{pmatrix},$$

and

$$\boldsymbol{\lambda}_g=\begin{pmatrix}\kappa_g\cos\omega\theta_g\\ \kappa_g\sin\omega\theta_g\end{pmatrix}.$$

The joint full conditional can be rewritten as:

(4)

$$P^*(A_{gj},\phi_{gj}) \propto \exp\left\{-\frac{1}{2\sigma_{gj}^2}\left(\boldsymbol{Y}'_{gj}-\boldsymbol{X}_j\boldsymbol{\beta}_{gj}\right)^T\left(\boldsymbol{Y}'_{gj}-\boldsymbol{X}_j\boldsymbol{\beta}_{gj}\right)+\boldsymbol{\beta}_{gj}^T\boldsymbol{\lambda}_g\right\}$$
$$\times\exp\left\{-\frac{1}{2\sigma_A^2}\left(A_{gj}-\mu_A\right)^2\right\}$$

And the conditional posterior density of $\boldsymbol{\beta_{gj}}$ is:

$$
\begin{aligned}
P^*\left(\boldsymbol{\beta}_{gj}\right) &\propto \exp\left\{-\frac{1}{2\sigma_{gj}^2}\left(\boldsymbol{Y}'_{gj}-\boldsymbol{X}_j\boldsymbol{\beta}_{gj}\right)^T\left(\boldsymbol{Y}'_{gj}-\boldsymbol{X}_j\boldsymbol{\beta}_{gj}\right)+\boldsymbol{\beta}_{gj}^T\boldsymbol{\lambda}_g\right\} \\
&\quad\times\frac{1}{\|\boldsymbol{\beta}_{gj}\|_2}\exp\left\{-\frac{1}{\sigma_A^2}\left(\|\boldsymbol{\beta}_{gj}\|_2-\mu_A\right)^2\right\} \\
&\propto \mathrm{BVN}\left(\left(\boldsymbol{X}_j^T\boldsymbol{X}_j\right)^{-1}\left(\boldsymbol{X}_j^T\boldsymbol{Y}_{gj}+\boldsymbol{\lambda}\sigma_{gj}^2\right),\left(\boldsymbol{X}_j^T\boldsymbol{X}_j\right)^{-1}\sigma_{gj}^2\right) \\
&\quad\times\frac{1}{\|\boldsymbol{\beta}_{gj}\|_2}\exp\left\{-\frac{1}{\sigma_A^2}\left(\|\boldsymbol{\beta}_{gj}\|_2-\mu_A\right)^2\right\}
\end{aligned} \tag{5}
$$

The slice sampling can then be performed similarly as Step II: Update within $\mathfrak{M}_1$: slice sampling for $A$ and $\phi$ in BayCT model.

**Step III: Updating between $\mathfrak{M}_1$ and $\mathfrak{M}_0$ using reversible jump sampling**

The parameter space of the two models that we want to select from are $\mathfrak{M}_0 : \boldsymbol{\beta_{0j}} = \left\{M'_{gj}, \sigma^{2\prime}_{gj}\right\}$ and $\mathfrak{M}_1 : \boldsymbol{\beta_{1j}} = \left\{M_{gj}, \sigma^2_{gj}, A_{gj}, \phi_{gj}\right\}$. The way to calculate the acceptance ratio of reversible jump and the sampling procedure are the same as Step III in BayCT.

**Step IV: Update $\Delta t_i$ in BayCT**

The full conditional of $\Delta t_i$ in the BayCT-M model is:

(6)

$$
\begin{aligned}
&P^*\left(\Delta t_i\right) \propto \prod_{gj:c_{gj}=1} P(Y_{gji}|A_{gj},\phi_{gj},t_i,\Delta t_i,M_{gj},\sigma^2_{gj})\cdot P\left(\Delta t_i|\theta',\kappa'\right) \\
&\propto \exp\left\{\sum_{gj:c_{gj}=1} -\frac{1}{2\sigma^2_{gj}}\left[Y_{gji}-M_{gj}-A_{gj}\cos\left(\omega(t_i+\Delta t_i-\phi_{gj})\right)\right]^2+\kappa'\cos\left(\omega(\Delta t_i-\theta')\right)\right\},
\end{aligned}
$$

where $\theta' = 0$ and $\kappa' = \kappa_t$ as specified in prior.

Here, the set of genes with index $c_{gj} = 1$ is chosen to update $\Delta t$. Without prior knowledge, we update $gj : c_{gj} = 1$ as the rhythmic genes identified in the last MCMC iteration, i.e., $\{gj : c_{gj} = 1\} = \{gj : \rho_{gj} = 1\}$. We can also pre-select a small set of core clock genes with known strong circadian rhythmicity, denoted by $s_g = 1$, and $\Delta t$ is updated with genes that are both pre-selected and estimated as rhythmic, i.e., $\{gj : c_g = 1\} = \{gj : s_g \cdot \rho_{gj} = 1\}$. Then, we can follow the same procedures as Section 1.1 to re-parameterize $P^*\left(\Delta t_i\right)$ and sample $\Delta t_i$

### 2. Supplementary Figures.

This section presents additional figures.

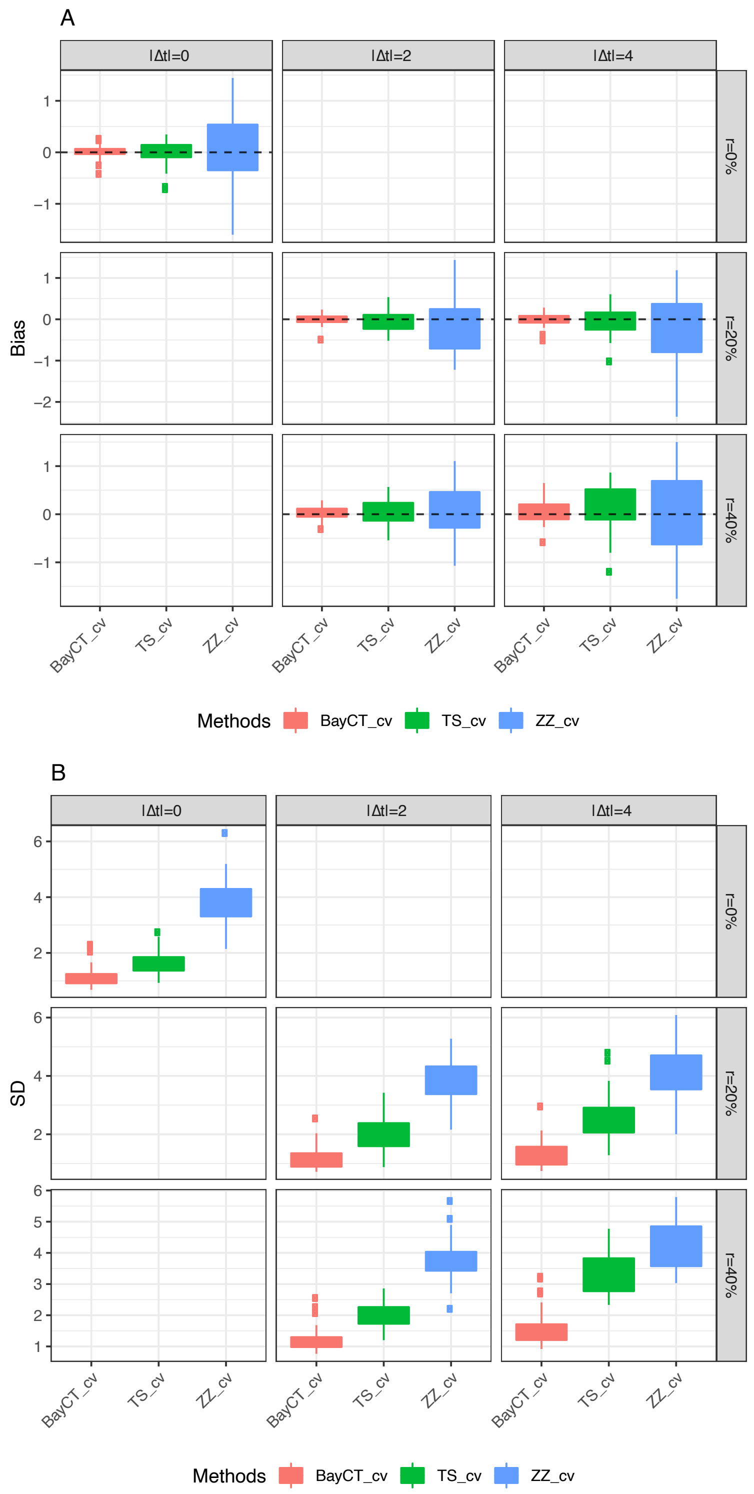


FIG S1. *Bias and standard deviation (SD) of the MCT estimations with cross-validation.*

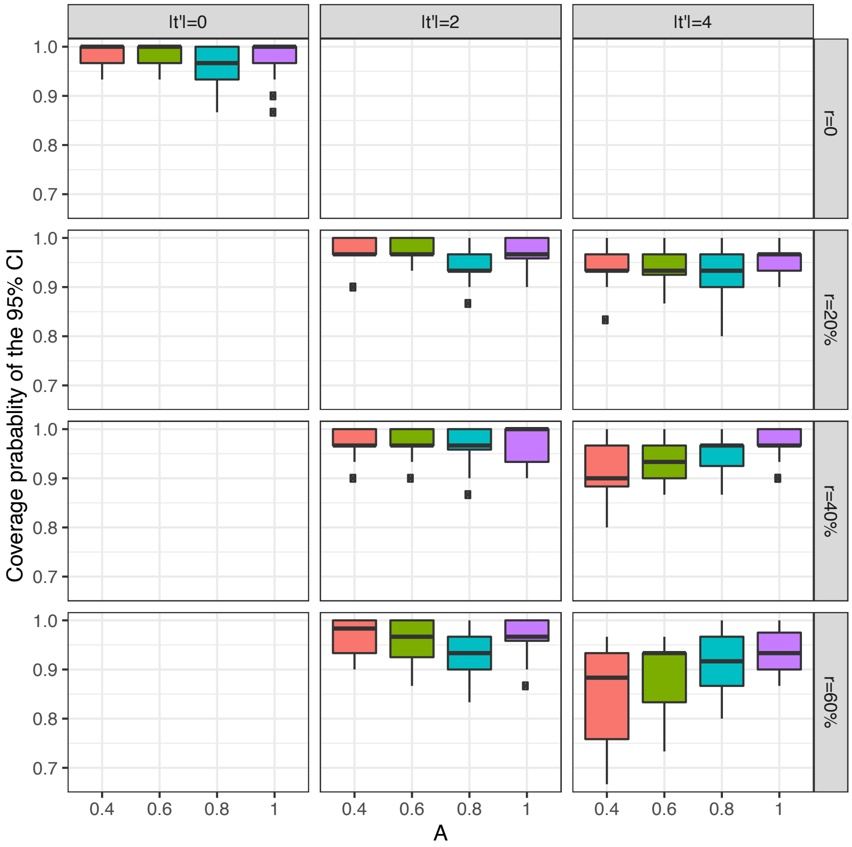


FIG S2. *Coverage rate of the 95% credible interval of the true MCT value.*

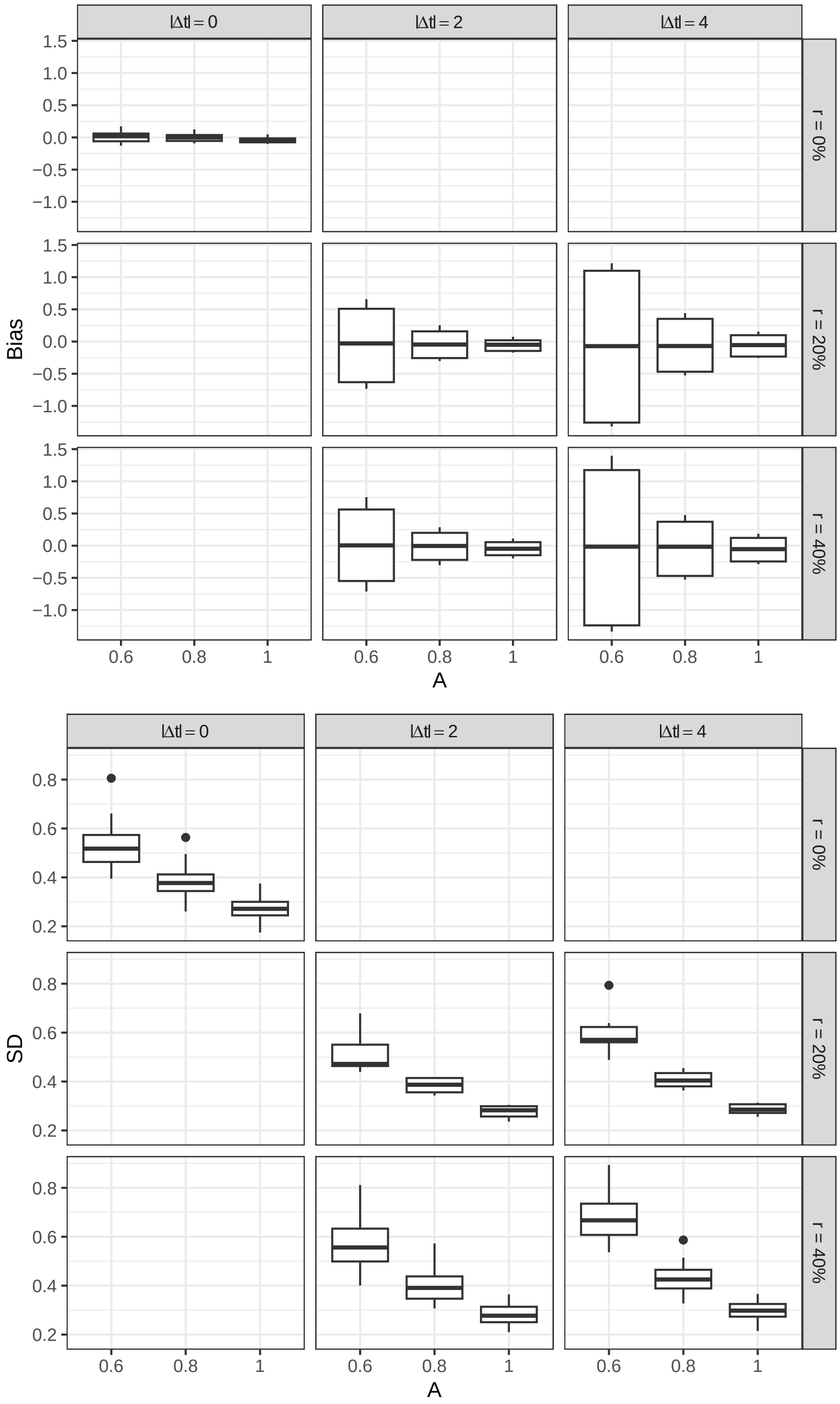


FIG S3. *Bias and standard deviation (SD) of the MCT estimations for BayCT-M SamePhase simulations with increasing amplitude A. The columns, moving from the left to right, represent an increasing magnitude of $\Delta t$, while the rows, from top to bottom, present an increasing percentage of samples affected by $\Delta t$.*

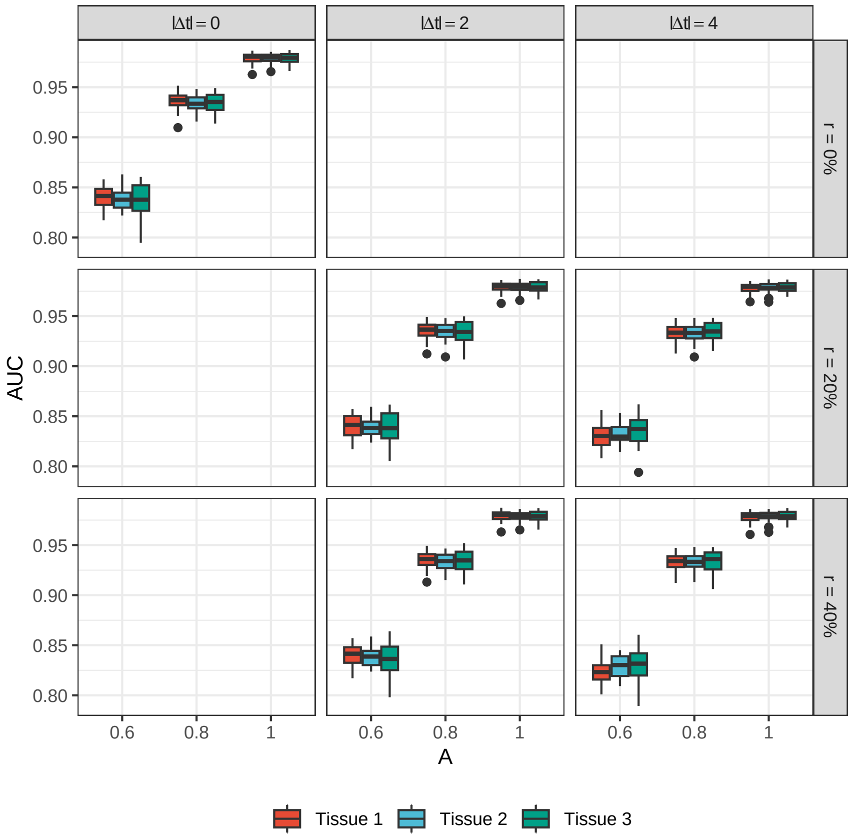


FIG S4. *AUC of the rhythmicity accuracy for BayCT-M SamePhase simulations with increasing amplitude A. The columns, moving from the left to right, represent an increasing magnitude of* $\Delta t$*, while the rows, from top to bottom, present an increasing percentage of samples affected by* $\Delta t$*.*

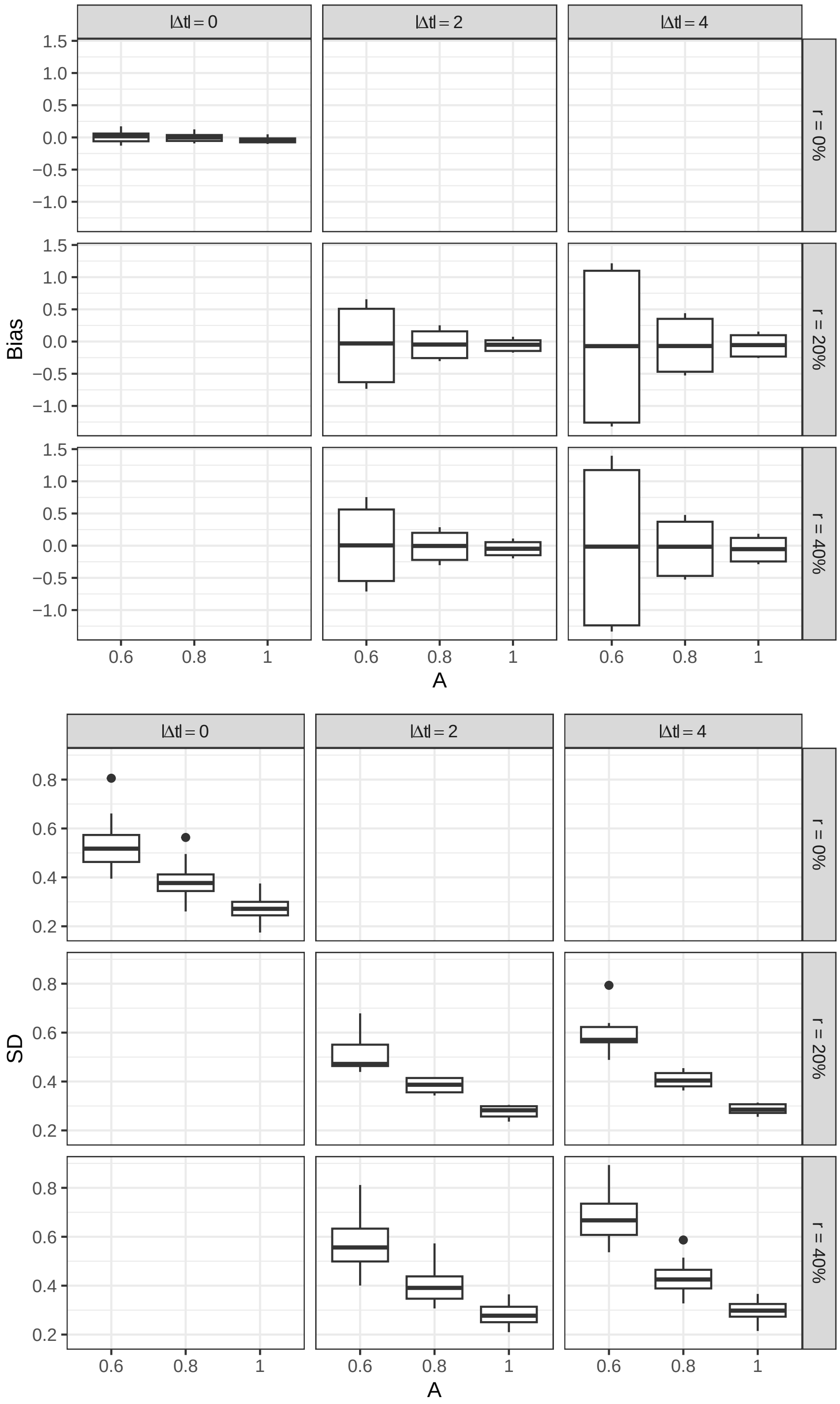


FIG S5. *Bias and standard deviation (SD) of the MCT estimations for BayCT-M DiffPhase simulations with increasing amplitude A. The columns, moving from the left to right, represent an increasing magnitude of* $\Delta t$*, while the rows, from top to bottom, present an increasing percentage of samples affected by* $\Delta t$*.*

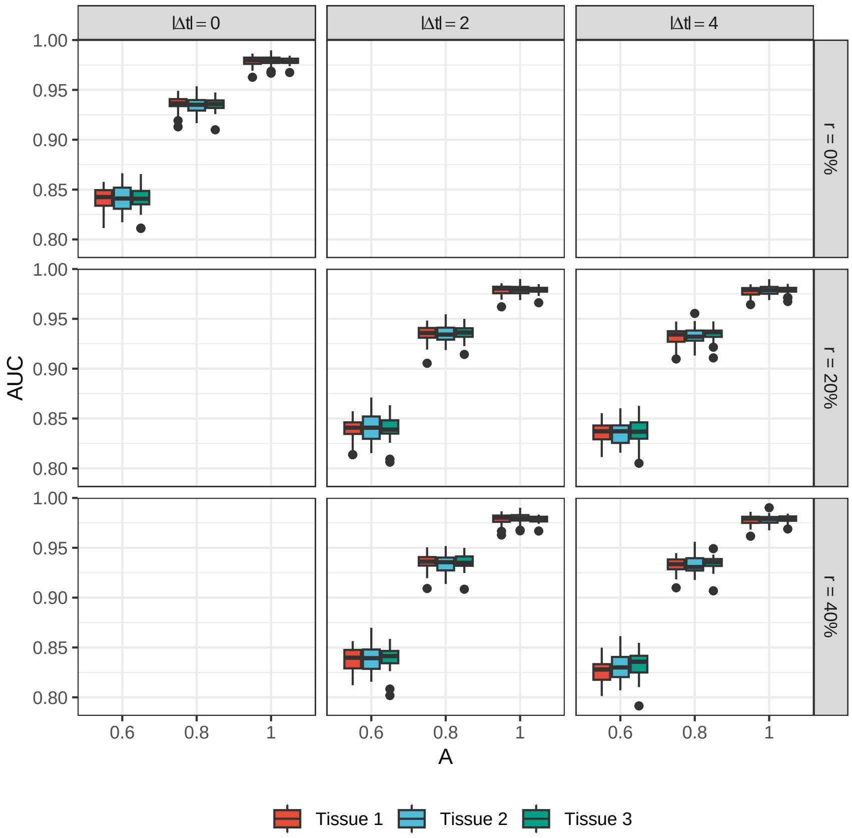


FIG S6. *AUC of the rhythmicity accuracy for BayCT-M DiffPhase simulations with increasing amplitude A. The columns, moving from the left to right, represent an increasing magnitude of $\Delta t$, while the rows, from top to bottom, present an increasing percentage of samples affected by $\Delta t$.*

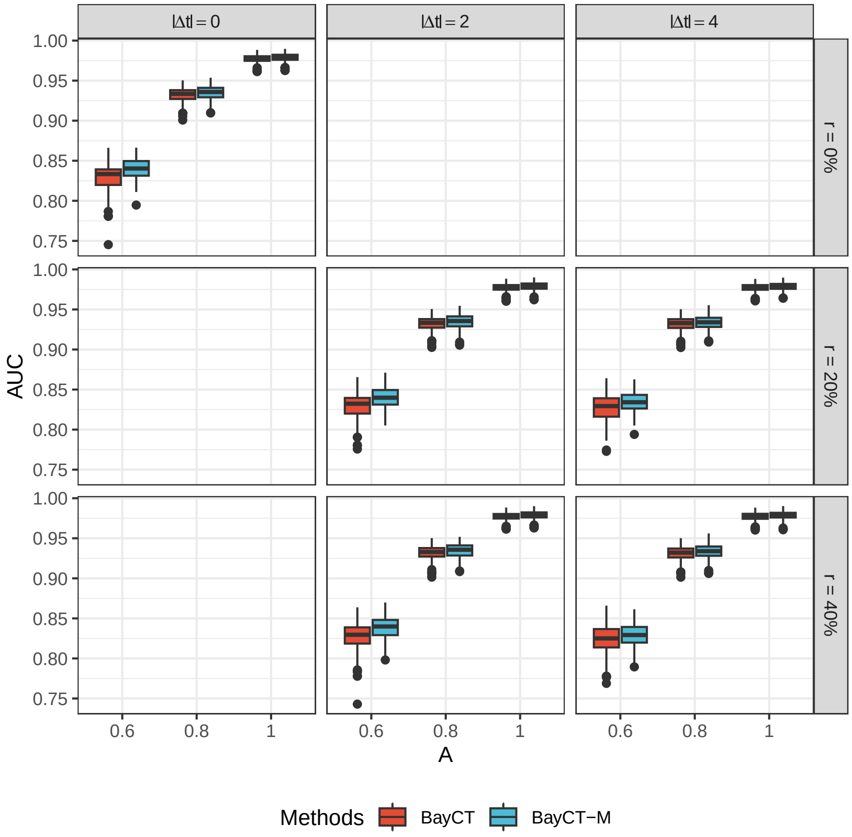


FIG S7. *Comparison of rhythmicity accuracy (AUC) between BayCT and BayCT-M. The columns, moving from the left to right, represent an increasing magnitude of $\Delta t$, while the rows, from top to bottom, present an increasing percentage of samples affected by $\Delta t$.*

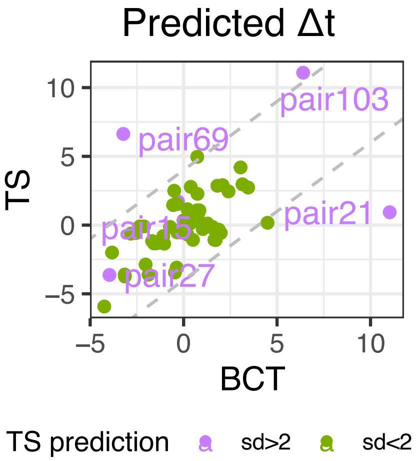


FIG S8. *Comparison of predicted $\Delta t$ from TimeSignature (TS) and BayCT (BCT). The predictions are colored with standard deviation of TS prediction of $\Delta t$ across three brain regions. The dashed lines have y-intercepts at* $4$*, and* $-4$.
